# Targeting temporal dynamics of microenvironmental factors halts tumor migration and alleviates effects of dynamic tumor heterogeneity

**DOI:** 10.1101/191221

**Authors:** Manjulata Singh, Xiao-Jun Tian, Vera S. Donnenberg, Alan M Watson, Jingyu Zhang, Laura P. Stabile, Simon C. Watkins, Jianhua Xing, Shilpa Sant

## Abstract

Targeting microenvironmental factors that foster migratory cell phenotypes is a promising strategy for halting tumor migration. However, lack of mechanistic understanding of the process impedes pharmaceutical drug development. Using a novel 3D microtumor model with tight control over tumor size, we recapitulated tumor size-induced hypoxic microenvironment and emergence of migratory phenotypes in epithelial T47D breast microtumors as well as those of patient-derived primary metastatic breast cancer cells, mesothelioma cells and lung cancer xenograft cells (PDX). The microtumor models from various patient-derived tumor cells and PDX cells revealed upregulation of tumor secretome, matrix metalloproteinase-9 (MMP9), fibronectin (FN), and soluble E-cadherin (sE-CAD) consistent with the clinically reported elevated levels of FN and MMP9 in the patient breast tumors compared to healthy mammary gland. We further showed that the tumor secretome induces migratory phenotype in non-hypoxic, non-migratory small microtumors. Subsequent mathematical model analysis identified a two-stage microtumor progression and migration mechanism, i.e., hypoxia induces migratory phenotype in the early initialization stage, which then becomes self-sustained through positive feedback loop established among the secretome. Both computational and experimental studies showed that inhibition of tumor secretome effectively halts microtumor migration despite tumor heterogeneity, while inhibition of the hypoxia is effective only within a time window and is compromised by tumor-to-tumor variation of the growth dynamics, supporting our notion that hypoxia initiates migratory phenotypes but does not sustain it. In summary, we show that targeting temporal dynamics of evolving microenvironments during tumor progression can halt and bypass major hurdle of tumor heterogeneity.

## Introduction

Metastases to distant organs remain the major cause of mortality in cancer patients. Metastasis begins when epithelial cells acquire migratory phenotype, break away from the primary tumor, navigate through extracellular matrix and extravasate into the circulation to home to the secondary organs. Lack of mechanistic understanding of this metastatic process remains a major challenge in identifying novel and effective treatment strategies for metastatic diseases. Current cancer treatments face grand challenges of overcoming tumor heterogeneity (Donnenberg and Donnenberg, 2015b) and associated drug resistance (Donnenberg et al., 2007), drug toxicity, drug-induced selection of more malignant cells, and tumor relapse (Easwaran et al., 2014; Huang, 2013; Koren and Bentires-Alj, 2015). Recent studies identify tumor microenvironment and associated tumor cell plasticity as important contributors to tumor heterogeneity and drug resistance (Fischer et al., 2015; Wang et al., 2014; Zheng et al., 2015). Hence, an increasing number of emerging cancer treatments target the tumor microenvironment (Brouxhon et al., 2013; Cox et al., 2016; Overall and Lopez-Otin, 2002). Unlike molecular therapies targeted to the tumor cells, those targeting the microenvironmental factors are less prone to adaptive changes in the tumor cells and thus, emergence of resistant cell populations (Cox et al., 2016). However, success of therapies targeting the tumor microenvironment remains mixed. For example, a hypoxia is known to activate several transcription factors (Gilkes and Semenza, 2013) and epigenetic modifications (Kao et al., 2016), thus induce cell phenotype changes such as conventional or partial epithelial-to-mesenchymal transition (EMT) that contributes to drug resistance and possibly tumor migration as well as metastasis (Fischer et al., 2015; Grigore et al., 2016; Kao et al., 2016; Lundgren et al., 2009; Nieto et al., 2016; Thiery, 2002; Tian et al., 2013; Zhang et al., 2014; Zheng et al., 2015). However, hypoxia-targeted therapies combined with current standard-of-care therapies (*e.g.,* chemotherapy or radiotherapy) showed mixed outcomes in several clinical trials (Mack et al., 2008; Patel and Sant, 2016; von Pawel et al., 2000; Williamson et al., 2005). The failure is not so unexpected since tumor cells can maintain their aggressive behavior in non-hypoxic conditions after they extravasate away from the tumor mass, and thus, are not affected by an anti-hypoxic therapy applied after tumor migration program is turned on. Therefore, an effective treatment strategy needs to consider the temporal dynamics of microenvironmental factors and other molecular species involved in the intricate regulatory network that initiates, maintains and promotes tumor progression (Donnenberg and Donnenberg, 2015b).

Tumor size is an important prognostic factor, which directly affects diffusion of nutrients, oxygen and other soluble factors, as well as cell-cell interactions, thus, affecting tumor biology and drug response. A typical *in vitro* three-dimensional (3D) culture conditions such as matrigel cultures or spinner flasks (Gong et al., 2015; Nyberg et al., 2005; Sant and Johnston, 2017) generates a broad size distribution of microtumors, which presents a technical difficulty for systematically studying the effect of tumor size, and size-associated physicochemical changes in the tumor microenvironment. We adopted our microfabricated hydrogel microwell platform to generate hundreds of uniformly sized microtumors of multiple cancer cell lines including breast (e.g., MCF7, T47D, BT474, MDA-MB-231 and HCC1187) and head and neck cancer cells (e.g., Cal33, Cal27, PE/CA-PJ-49, SCC9, and BICR56) (Singh et al., 2015; Singh et al., 2016). We demonstrated that precise control over microtumor size is translated in controlled physicochemical microenvironments that also recapitulate inter- and intra-tumor heterogeneity (Singh et al., 2015; Singh et al., 2016; Singh et al., 2017). By manipulating tumor size alone without any other culture or genetic manipulations, we reproducibly generated two distinct phenotypes from the same non-invasive breast cancer cell line; small microtumors represent non-hypoxic and non-migratory phenotype whereas large microtumors exhibit hypoxic microenvironments with migratory phenotype (Singh et al., 2015; Singh et al., 2016; Singh et al., 2017). This observation was extended across multiple luminal breast cancer cell lines (*e. g.* MCF7 and T47D). Interestingly, we observed that the migratory phenotype in large microtumors was irreversible as they maintained their migratory phenotype even after dissociating them into single cell suspension and re-growing into small non-hypoxic microtumors (Singh et al., 2016). Therefore, our microtumor model system recapitulates both, hypoxia-induced migratory cell phenotype and its subsequent maintenance under hypoxia-free microenvironment.

In this study, we integrated mathematical modeling and *in vitro* experimental approaches using our 3D size-controlled microtumor models to test a two-stage tumor progression hypothesis that hypoxia induces migratory phenotype in the early initialization stage, which is subsequently maintained by other factors secreted in the hypoxic tumor microenvironment. We provide compelling evidence that targeting molecules involved in the maintenance stage, but not in the initialization axis of tumor progression, can be an effective treatment strategy that is less affected by tumor heterogeneity and tumor progression stage.

## Results

### Large microtumors exhibit hypoxia, collective migration and aggressive phenotype

Here, we adopted our microfabricated hydrogel microwell platform (Singh et al., 2015; Singh et al., 2016) to generate uniform size T47D microtumors of about 150 μm (referred as ‘small’ microtumors) and 500-600 μm in diameter (referred as ‘large’ microtumors). After seeding into the microwells, cells compact forming uniform size microtumors within 24 h and continue to grow over 6 days in culture (Singh et al., 2017). Each 2 × 2 cm^2^ hydrogel microwell array fabricated of non-adhesive polyethylene glycol dimethacrylate (PEGDMA) can generate approximately 1500-2100 small microtumors or 270-290 large microtumors. Each 6-day old small (150 μm) microtumor contains approximately 300-500 cells while large (600 μm) ones contain 6000-8000 cells per microtumor (Singh et al., 2017). This 3D microtumor platform allows for controlled and reproducible studies on initial avascular tumor growth and how tumor size-mediated microenvironmental changes affect tumor progression (Singh et al., 2016).

We cultured small and large size-controlled microtumors of non-invasive human T47D cells in microwell arrays for six days (**Fig. 1A**). We observed that cells seeded in the microwell compact during the first 24h, and continue to grow over six days in culture. While small microtumors remained non-migratory even after six days in culture, large microtumors moved towards the wall of the microwells and showed collective migration out of the wells (**Fig. 1A**) starting as early as day 3. We ruled out the differences in the proliferation or crowding as the cause of migration since we have already shown that both small and large microtumors exhibited similar increase in the cell number (about 1.5-2 fold) over a period of six days (Singh et al., 2017). We quantified the distance of migration (d), area of the migrated portion (X), and total microtumor area (Y) using ImageJ (**Fig. 1B**). Despite highly homogeneous microtumor size distribution, we observed microtumor-to-microtumor heterogeneity in the migration dynamics of large (600 μm) microtumors, parallel to the well-documented cell-to-cell heterogeneity (De Sousa et al., 2013). Indeed, these individual large microtumors displayed variations in the kinetics of migration, as quantified by their migrated distance (d) and the extent of migration (X/Y*100) (**Fig. 1C-D**). We conjectured that the different behaviors of 150 and 600 μm microtumors result from size-associated changes in their microenvironments. Therefore, we measured intra-tumoral expression of hypoxia-inducible factor 1 alpha (HIF-1α), EMT markers (vimentin, VIM and fibronectin, FN) and levels of tumor secretome including matrix metalloproteinases (MMPs), FN and soluble E-cadherin (sE-CAD) secreted in the conditioned media (CM). Protein expression results revealed upregulation of HIF-1α in tumor cell lysate of 600 μm microtumors **(Fig. 1E)**. We also observed upregulation of mesenchymal markers, VIM and FN without loss of epithelial marker, E-CAD (**Fig. 1F**) suggesting that the migratory behavior of large microtumors may not be due to complete EMT; instead it represents “cohesive or collective migration” that requires E-CAD (Cheung and Ewald, 2014). Indeed, co-staining of E-CAD and VIM in large T47D microtumors showed heterogeneous population of cells co-expressing E-CAD and VIM (**Fig. 1G**, white arrows in bottom panel) along with the epithelial cells expressing only E-CAD (**Fig. 1G**, green). ELISA measurement of sE-CAD and western blot of conditioned media (CM) showed upregulation of FN and sE-CAD while gelatin zymography displayed increased levels of Pro-MMP9 and MMP9 in the 600 μm microtumors compared to the 150 μm ones (**Fig. 1H-I**). We also observed activation of the downstream extracellular signal-regulated kinase (ERK) as shown by upregulation of pERK in large microtumors (**Fig. 1J**). These results indicate that increased tumor size induces hypoxic tumor microenvironments, which further promote collective migration in large microtumors possibly due to the acquisition of mesenchymal features without loss of E-CAD. In addition, large microtumors also exhibit intra-tumoral heterogeneity in the expression of E-CAD/VIM and inter-tumoral heterogeneity in migration kinetics along with increased levels of tumor secretome (sE-CAD, FN, MMP9).

**Figure 1:**
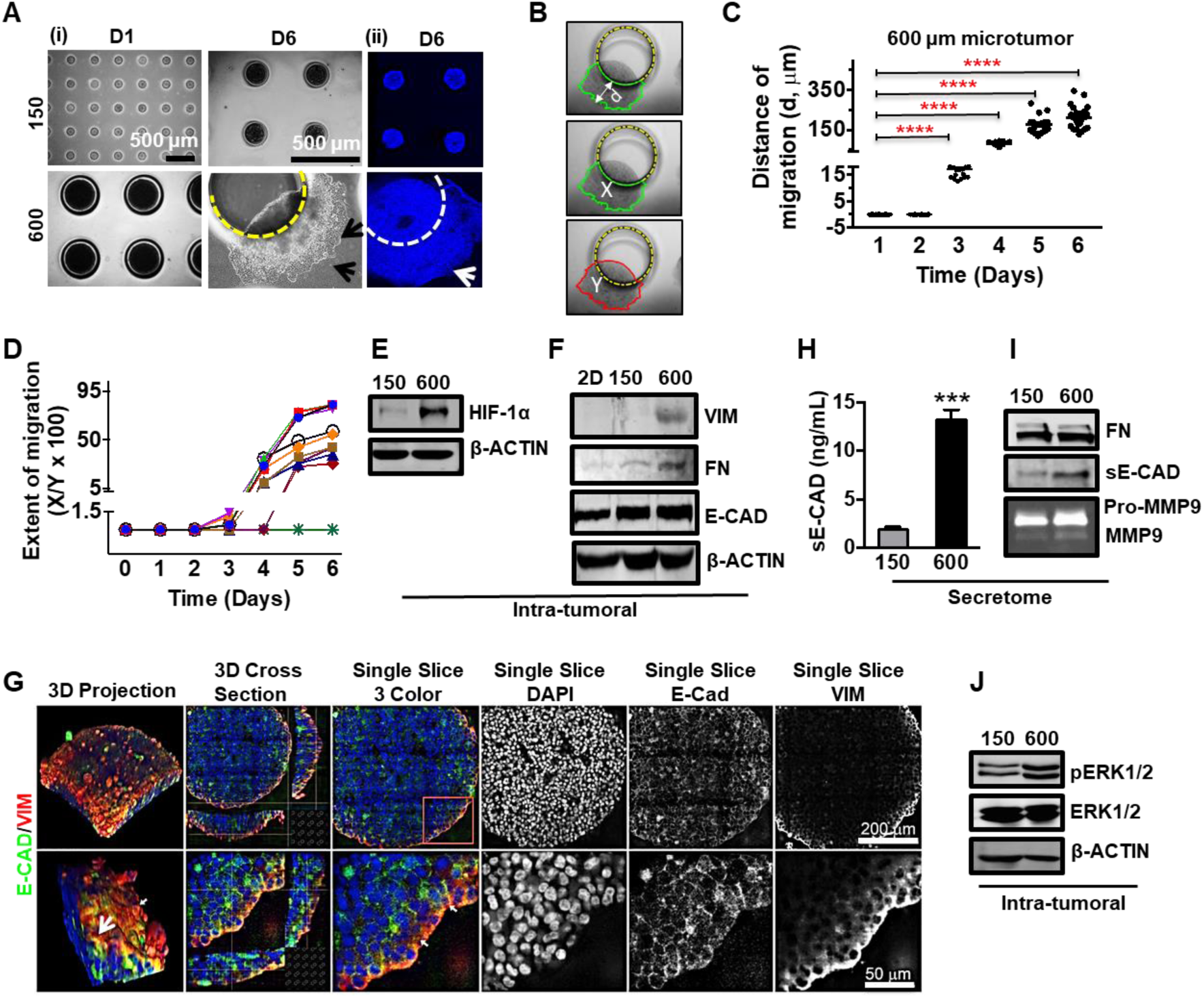
Large microtumors exhibit hypoxia, collective migration and aggressive phenotype. Size-controlled 150 and 600 μm microtumors of T47D cells were fabricated using PEGDMA hydrogel microwell arrays. (A-i) Photomicrographs showing microtumors on day 1 and day 6. Large microtumors (600 μm) showed migratory phenotype on day 6 (black arrows); (A-ii) Hoechst-stained 600 μm microtumor showing migratory phenotype (white arrow) on day 6; (B) Schematics defining distance of migration (d), area of microtumor migrated out of the microwell (X) and total area of the microtumor (Y); (C) Distance of migration for individual microtumors was estimated each day by measuring more than 10 perpendicular distances at migratory front from the periphery of microwell and averaging them. Statistical analysis was done by unpaired t-test by comparing the mean distance of each day with day 1. Mean migration distances significantly increased from day 1 to day 6 with significance of p****< 0.0001 w.r.t. day 1. Each dot represents data for individual microtumor. Please note the broad distribution highlighting inter-tumoral heterogeneity in migration distance, especially on day 5 and 6. (D) Migration trajectories of individual 600 μm microtumors from day 1 to day 6 showing inter-tumoral heterogeneity in migration kinetics; Extent of migration for each time point was calculated as migrated distance/the total area of microtumor X 100. Each line represents migration kinetics for individual microtumor. (E) Large (600 μm) microtumors exhibited upregulation of HIF-1α expression (intra-tumoral) and (F) high levels of VIM and FN expression without significant change in E-CAD protein levels; (G) Z-stacked images of large microtumors showed simultaneous expression of E-CAD (green) and VIM (red) (white arrow) demonstrating co-expression of E-CAD and VIM on the periphery of the microtumor while maintaining E-CAD expression. Grayscale images are 2D sections through depth of 600 μm microtumor and showed the expression of E- CAD (throughout the tumor) and VIM (only on periphery); (H) Large (600 μm) microtumors showed significantly higher expression of sE-CAD in hypoxic secretome (conditioned media, CM) measured by ELISA (***p<0.001 w.r.t 150 μm microtumors, unpaired t-test) and western blot (I) and showed higher expression of FN in CM as compared to 150 μm microtumors. Similarly, gelatin zymography showed higher levels of Pro-MMP9 and MMP9 in 600/CM as compared to 150/CM; (J) Large microtumors also showed higher pERK expression compared to small microtumors. Data presented is representative from three independent experiments (n=3).

### Non-hypoxic, non-migratory small microtumors exhibit collective migration upon treatment with recombinant human sE-CAD (Rec-sE-CAD) and hypoxic secretome containing sE-CAD

Next, we asked whether hypoxia alone is responsible for inducing the migratory phenotype in large microtumors. In previous work, HIF-1α inhibition either by siRNA or chemical inhibitor treatments suppressed migration of large microtumors only partially (Singh et al., 2016). Moreover, when non-hypoxic small microtumors were dissociated into single cells and re-grown into large ones, the cells acquired migratory phenotype, confirming the role of the hypoxic microenvironment in the process (Singh et al., 2016). Interestingly, when large microtumors were dissociated into single cells to remove the hypoxic microenvironment and re-grown into small microtumors, they maintained their migratory phenotype (**Fig. 2A**). This suggests that the hypoxia-induced migratory phenotype is not reversible even after removal of the hypoxic microenvironment. We hypothesized that the microtumor secretome contains some of the self-maintenance factors. To test this hypothesis, we treated non-hypoxic, non-migratory 150 μm microtumors with the hypoxic secretome (CM) of 600 μm microtumors (600/CM) from day 3 until day 6. 600/CM treatment induced the migration in more than 60% of otherwise non-migratory 150 μm microtumors (**Fig. 2B-C**).

**Figure 2.**
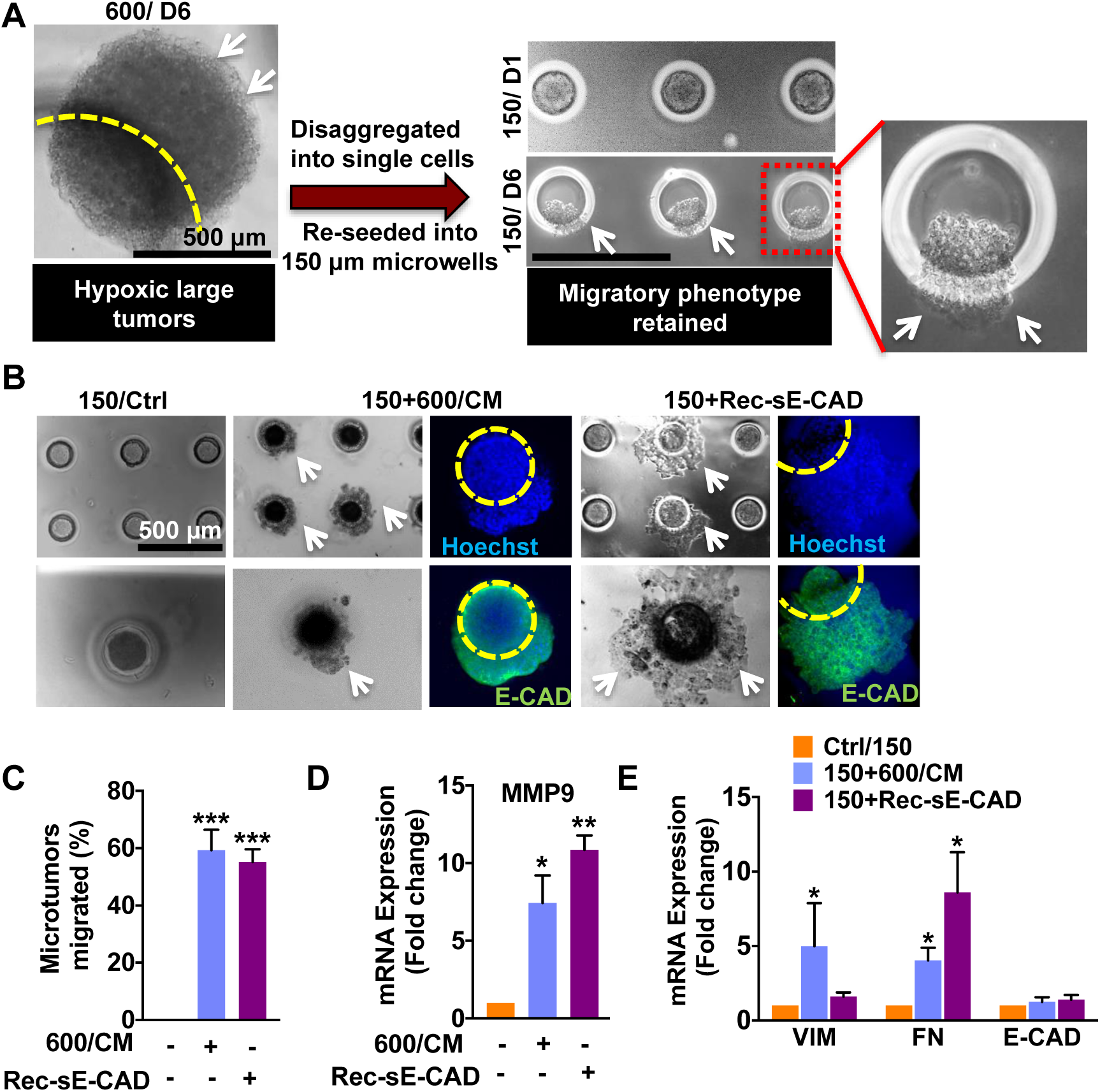
Recombinant sE-CAD and hypoxic secretome induce collective migration in non-migratory 150 μm microtumors. To confirm the plasticity of migratory phenotype, 6-day old 600 μm microtumors were disaggregated and re-seeded into 150 μm hydrogel microwell devices. (A) 600 μm microtumors maintained their migratory phenotype (white arrows) even after disrupting hypoxia (disaggregation) and re-growing them into non-hypoxic 150 μm sizes (Inset). To investigate the role of hypoxic secretome and sE-CAD in regulating migratory phenotype, 150 μm microtumors were fed either with the conditioned media of 600 (600/CM) or with 20 μg/mL recombinant sE-CAD (Rec-sE-CAD). (B) Representative images showing collective migration (white arrows) in 600/CM and Rec-sE-CAD treated 150 μm microtumors without loss of E-CAD (green), cell nuclei are stained with Hoechst (blue) and yellow circle marks the boundary of a microwell; (C) Quantification of migration showed more than 60% migration for 150 μm microtumors post treatments. Data are presented as mean ±SD from at least three independent experiments; ***p ≤ 0.001 w.r.t. untreated (control) 150 μm microtumors; (D) Both Rec-sE-CAD and 600/CM treatments upregulated mRNA expression of *MMP9;* and (E) *FN* in treated 150 μm microtumors compared to untreated control; however, *VIM* was upregulated only in Rec-sE-CAD treated microtumors. Data are presented as mean ±SEM from three independent experiments. *p<0.05, **p<0.01, ***p<0.001 w.r.t. untreated control (unpaired t-test).

The microtumor secretome contains multiple factors, and sE-CAD is one of them that we observed in higher amounts in the 600/CM than that in the 150/CM (**Fig. 1H-I**). Treatment with recombinant human sE-CAD (Rec-sE-CAD, Fc chimera) also induced migration in more than 60% small microtumors (**Fig. 2B-C**). Both, 600/CM and Rec-sE-CAD treatments induced collective migration without loss of E-CAD (green, **Fig. 2B**). sE-CAD is known to upregulate MMP expression (Brouxhon et al., 2014b). To investigate the effect of 600/CM and Rec-sE-CAD on MMP expression, we measured mRNA expression of *MMP9* in the treated microtumors. Quantitative PCR results showed more than 8-fold increase in *MMP9* expression in both 600/CM and Rec-sE-CAD treated 150 μm microtumors (**Fig. 2D**). To further characterize the changes in cell phenotype in small microtumors treated with either 600/CM or Rec-sE-CAD, the mRNA expression of selected epithelial and mesenchymal markers was measured in treated microtumors, and compared with untreated small microtumors (controls). The treatment with 600/CM significantly upregulated *VIM* (5.0± 2.9 folds) and *FN* (4.0± 0.9 folds) compared to 150/control, while sE-CAD (Rec-sE-CAD) treatment significantly upregulated *FN* (8.6± 2.7 folds) with no effect on *VIM* **(Fig. 2E)**. These results confirmed that either hypoxic secretome (600/CM) or sE-CAD alone could induce the migratory phenotype in non-migratory (150 μm) epithelial microtumors although they have differential effects on regulating mesenchymal markers such as *VIM*. In comparison, CM of 150 μm microtumors had no effect on the migratory behavior of either 150 or 600 μm microtumors **(Supplementary Fig. 1).**

### Tumor secretome in patient tumors, microtumors of primary patient-derived tumor cells and PDX cells showed upregulation of FN, MMP9 and sE-CAD

To investigate the clinical relevance of tumor secretome, FN and MMP9, we extracted clinical data from various breast cancer types from a set of 3735 human samples by Genevestigator® and examined them with AffyU133Plus2. Expressions of FN and MMP9 were higher in various breast carcinomas as compared to the normal mammary gland (**Fig. 3A**). Furthermore, we fabricated small and large microtumors using primary cancer cells derived from fresh metastatic breast cancer (mBC, one patient sample), mesothelioma (one patient sample) and lung PDX-derived cells (two different patient samples) (**Fig. 3B**). Despite the metastatic and highly invasive nature of malignant pleural effusions, morphological characterization of mBC cells revealed microtumor clusters (**Supplementary Fig. 2A)**. Flow cytometry characterization of flow sorted aneuploidy tumor cells revealed mixed population of cytokeratin (CTK)+/VIM+ and only VIM+ cells with small subpopulation of cells expressing surface mesenchymal markers (CD44, CD90). These patient- and xenograft-derived primary cancer cells formed small and large microtumors in our hydrogel microwell arrays, albeit, the initial microtumor sizes right after seeding were smaller than breast cancer cell line-derived microtumors (**Supplementary Fig. 3A, B and C**). This could be attributed to the differences in the cell size as well as compaction ability of individual cells isolated from different tumors. We have reported similar observations with head and neck cancer cell lines earlier (Singh et al., 2015; Singh et al., 2016). Notably, the microtumors of primary cancer cells obtained from different tumor types also recapitulated the results obtained in the breast cancer cell lines, with large microtumors showing migratory phenotype on day 6-8 while smaller ones remaining non-migratory. Compared to the small microtumors, the large microtumors of patient-derived mBC cells showed 4-fold upregulation of mesenchymal marker *FN* with a slight but not significant change in *VIM* (**Fig. 3C**). Interestingly, we also observed significant downregulation of *E-CAD* and *MMP9* (**Fig. 3C, D**). This could be because the primary mBC cells are derived from highly metastatic pleural effusions and may have molecular subtype could be different than T47D cells. Despite the microtumor clusters present in mBC pleural effusions as shown here, the mesenchymal phenotype is predominant (Donnenberg et al., 2013). We also observed increased concentration of sE-CAD (5.56 ±0.77 ng/mL) in 600 μm as compared to 2.1 ±0.32 ng/mL in 150 μm microtumors measured by ELISA (**Fig. 3E**). Western blot revealed upregulation of FN and sE-CAD in conditioned media (secretome) collected from large mBC microtumors compared to small non-migratory microtumors (**Fig. 3F)**. Similarly, we observed significant upregulation of *FN* (2.5-fold) and *MMP9* (5-fold) mRNA in large microtumors of lung PDX compared to small microtumors **(dotted line, Fig. 3G and H)**. Secretome analysis by ELISA and western blots showed high levels of sE-CAD and FN in the conditioned media of large microtumors of PDX and mesothelioma (**Fig. 3I-J** and **Supplementary Fig. 3D-E)**. Small lung PDX microtumors had non-detectable levels of sE-CAD measured by ELISA. These results indicate that our microtumor model can be easily adapted to primary cells derived from patient tumors and from human xenografts across multiple tumor types such as breast, mesothelioma and lung tumors.

**Figure 3.**
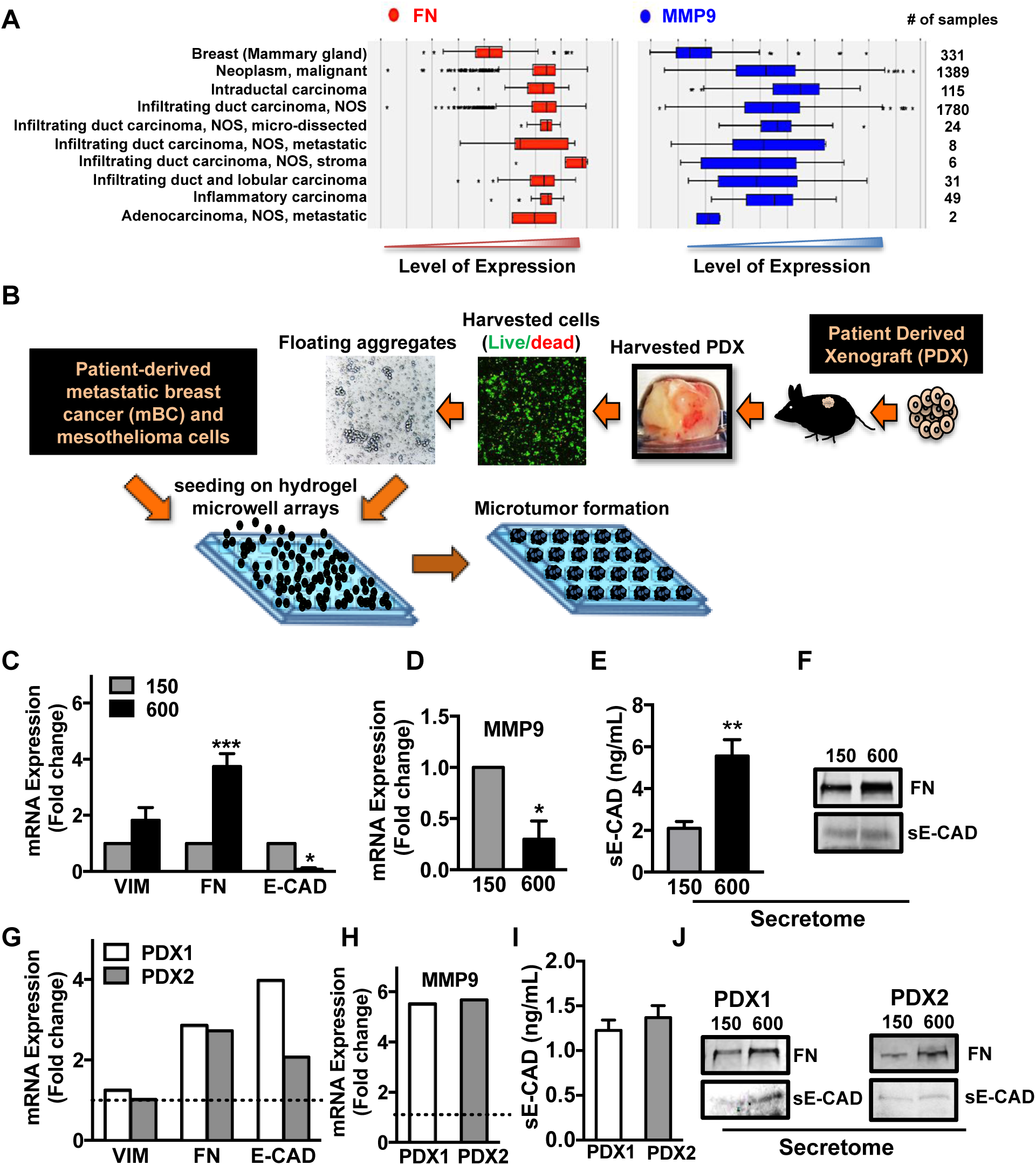
Tumor secretome in patient tumors, microtumors of primary tumors and PDX cells showed upregulation of FN, MMP9 and sE-CAD. (A) To establish clinical correlation of the T47D microtumor results with the patient data, clinical data set was extracted for FN and MMP9 status. The results showed higher levels of FN and MMP9 in breast carcinomas as compared to normal mammary gland. (B) Hydrogel microwell arrays (150 and 600 μm) were used to generate the microtumors from patient-derived primary metastatic breast cancer (mBC) cells (1 patient), mesothelioma cells (1 patient) and from non-small cell lung cancer (NSCLC) PDX model established using lung cancer metastasized to brain (2 PDX tumors). Schematic shows the method of microtumor fabrication using patient-derived primary metastatic breast cancer, mesothelioma and mouse lung PDX cells; (C-F) mBC cell-derived microtumors were cultured up to 8 days and harvested for mRNA, ELISA and protein analysis. (C) qRT-PCR results showed upregulation of *FN* and downregulation of *E-CAD* and (D) M*MP9* mRNA expression in large mBC microtumors as compared to the small ones; *p<0.05, ***p<0.001 w.r.t. 150 μm microtumors, unpaired t-test. (E) ELISA showed higher sE-CAD levels and (F) Western blots with the conditioned media (CM) showed increased FN in large mBC microtumors compared to the small ones; (G-J) PDX cell-derived microtumors were cultured up to 8 days and harvested for mRNA, ELISA and protein analysis. (G) Large microtumors of both PDX samples showed upregulation *FN*, *E-CAD* and (H) *MMP9* mRNA expression compared to the small 150 μm microtumors (dotted line in the graph); (I) Large PDX microtumors showed higher sE-CAD levels compared to non-detectable levels in small microtumors by ELISA; (J) Western blots from CM also showed upregulation of FN and sE-CAD in the secretome of large PDX microtumors suggesting the role of microtumor size in altering the tumor secretome and subsequently the tumor phenotype.

### Mathematical model predicts a two-stage mechanism for microtumor progression and migration

Based on the above experimental findings in T47D microtumors, our previous study (Singh et al., 2016), we constructed a minimal regulatory network describing how migration of a microtumor is coupled to its microenvironment (**Fig. 4A and Supplementary Fig. 4**). Increase in microtumor size leads to hypoxia, which stabilizes HIF-1α. HIF-1α can increase the expression of MMP9 (Choi et al., 2011), one of the proteases that can generate sE-CAD by promoting cleavage of E-CAD (Maretzky et al., 2005). Abnormally elevated levels of sE-CAD have been reported in various types of cancers as shown by clinical data and published reports (David and Rajasekaran, 2012; De Wever et al., 2007; Repetto et al., 2014). sE-CAD can bind and activate the epidermal growth factor receptor (EGFR) family (HER1-4), which further activates the ERK pathway (Brouxhon et al., 2014c; Inge et al., 2011; Najy et al., 2008) involved in the tumor cell proliferation. Moreover, pro-MMP9 and FN are targets of ERK-1/2 (Brouxhon et al., 2014c; Grabowska et al., 2012; Moshal et al., 2006; Overall and Lopez-Otin, 2002). For simplicity, these intermediate regulators (pro-MMP9, ERK-1/2) are taken implicitly in our minimal model. This completes a positive feedback loop formed among the microtumor secretome species, including MMP9, sE-CAD and FN. We hypothesized that this positive feedback is critical for collective migration of microtumors. Thus, we built a mathematical model to describe the dynamics of these regulators with a set of ordinary differential equations (ODEs) as shown in **Fig. 4B** (see Supplementary material for details).

**Figure 4.**
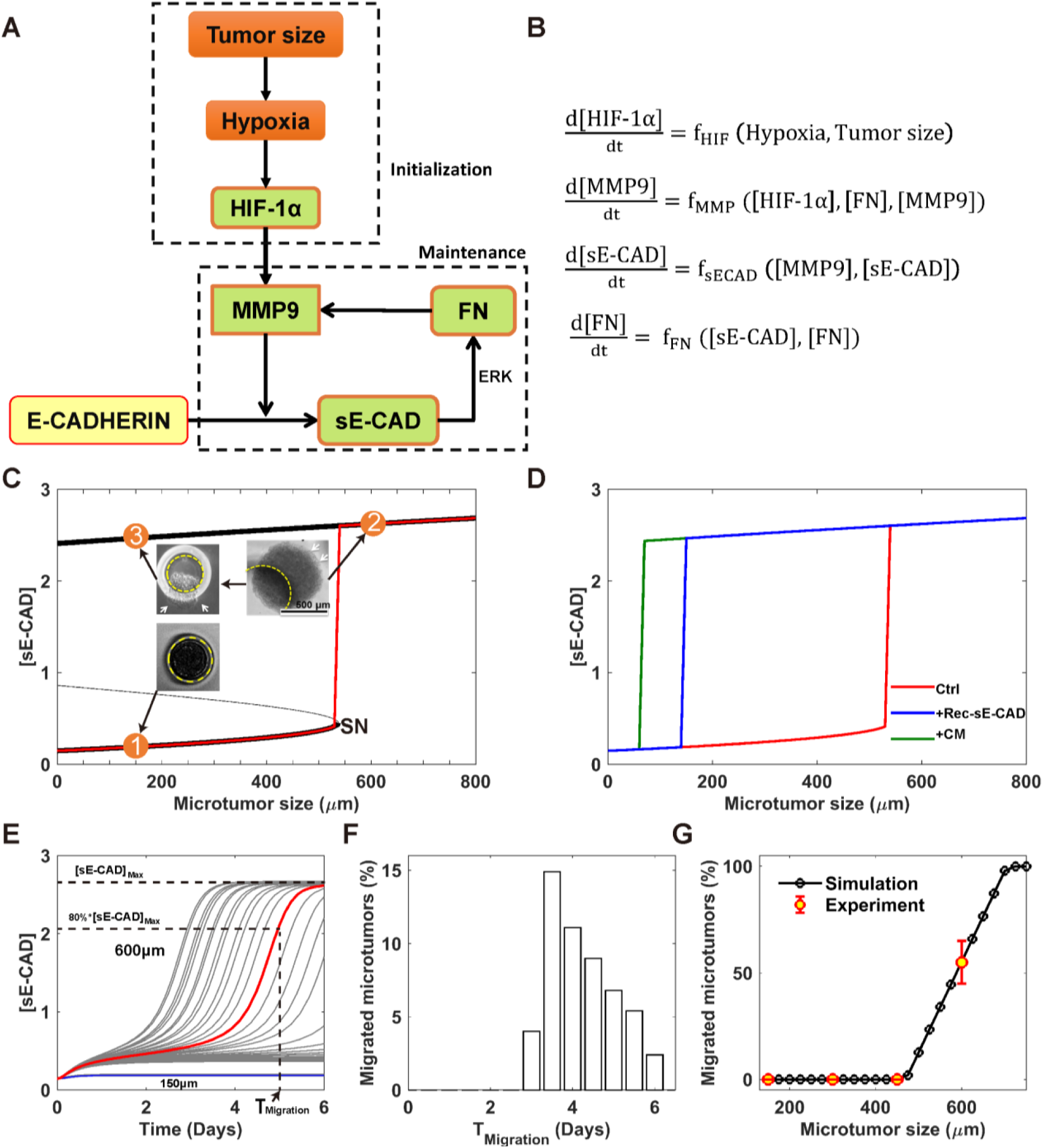
Mathematical model predicts an irreversible bistable switch as the mechanism of the migratory behavior of the microtumor. (A) Schematic depiction of the core regulatory network. HIF-1α is stabilized by hypoxia, which is induced with the increase in the microtumor size. Once HIF-1α is accumulated, it increases the expression of MMP9. MMP9 are one of the proteases, which can increase the *γ*-secretase mediated cleavage of E-CAD and thus, generate sE-CAD. sE-CAD is able to induce the expression of FN through the activation of ERK pathway. Moreover, FN upregulates MMP9. That is, a positive feedback loop is formed among MMPs, sE-CAD and FN; (B) The corresponding mathematical model described with ordinary differential equations (ODE) (see a detailed description in the Supplemental Information); (C) Bifurcation analysis of the mathematical model gives an irreversible bistable switch. There is a threshold (SN) of microtumor size for the activation of this switch. As shown in case 1, 150 μm microtumor does not exceed the threshold and does not migrate; in case 2, hypoxic microenvironment in 600 μm microtumor exceeds the threshold and induces high levels of [sE-CAD] and a migratory phenotype. The irreversibility of the switch is very important for the collective migration since the microtumor can stay in the migratory state even after it escapes from the original site and migrates collectively like a sheet. As shown in case 3, re-growing 600 μm microtumors into non-hypoxic small microtumors still maintains high [sE-CAD] and migratory phenotype; (D) The threshold of microtumor size for migration shifts to lower sizes (< 200 μm) in the presence of recombinant sE-CAD (Rec-sE-CAD) or conditional media from migrated microtumor (CM); (E) The heterogeneous dynamics of the [sE-CAD] for 150 μm and 600 μm microtumors due to the inter-tumoral heterogeneity. Here, it is assumed that the microtumor migration becomes visible when [sE-CAD] exceeds 80% of its steady state value with the corresponding time, T_migration_. Thirty representative microtumors are shown for each 150 μm and 600 μm microtumors; however, unlike 600 μm microtumors, 150 μm microtumors have negligible heterogeneity; (F) The distribution of T_migration_ among microtumors with 1000 representative microtumors; (G) Both, simulation and experimental data reveals that the percentage of the migrated microtumor depends on the microtumor size.

Through bifurcation analysis, we found that within a broad parameter range, the mathematical model gives an irreversible bistable switch. The level of [sE-CAD] is low in the secretome of small microtumors (**Fig. 4C**, point 1). With increase in the tumor size, [sE-CAD] initially increases very slowly until a threshold (Saddle-Node bifurcation point, SN) of the microtumor size is reached, above which [sE-CAD] jumps to a higher value (**Fig. 4C,** point 2). Here we used [sE-CAD] as a marker since it is experimentally straightforward to measure and we have shown that the CM of migratory 600 μm microtumors contains higher [sE-CAD] than that of non-migratory 150 μm microtumors (**Fig. 1H-I**). It should be noted that this bistable switch is irreversible since [sE-CAD] remains high even after large microtumors are dissociated and re-grown as small ones **(Fig. 4C**, point 3), just as what we observed experimentally (**Fig. 2A**). The irreversibility of the switch is important for the collective migration since it implies that cells in the large microtumor can maintain their migratory phenotype even after escaping the hypoxic core or disaggregating from the tumor. This mathematical analysis suggests a two-stage tumor progression mechanism (**Fig. 4A**), where hypoxia is important in the early initiation stage of tumor progression, and components of the MMPs/sE-CAD/FN feedback loop work synergistically to maintain the migratory phenotype. Indeed, the modeling results in **Fig. 4D** predict that treatment of microtumors with 600/CM (containing MMP9 + sE-CAD, green line) or recombinant human sE-CAD (Rec-sE-CAD, blue line) shifts the threshold of microtumor size below 200 μm for [sE-CAD] and tumor migration propensity. This was confirmed by our experimental results where treatment with 600/CM or Rec-sE-CAD induced migration in otherwise non-migratory 150 μm microtumors (**Fig. 2B-C**). It is noted that patient-derived metastatic breast cancer (mBC) microtumors start to migrate before they reach the size of the 500-600 μm (**Supplementary Fig. 3)**. That is, the threshold of the migration for these metastatic microtumors is smaller than that of T47D microtumors. This may be due to the differences in the cell/tumor subtype, baseline phenotype of the cells, etc.

Our model studies reveal that a microtumor with size exceeding the critical value (SN) (**Fig. 4C**) generates and then maintains a microenvironment with high [sE-CAD] and starts to migrate. Due to stochasticity such as cell growth and division rates, the onset as well as the kinetics of individual microtumor migration may vary, which we call inter-tumor dynamic heterogeneity here. **Fig. 4E** shows simulated temporal dynamics of how [sE-CAD] in the CM of individual 150 and 600 μm microtumors increases with time, where the inter-tumor dynamic heterogeneity manifests itself, especially for the 600 μm microtumors. Consequently, individual microtumors start to migrate at different times after day 2 (**Fig. 4F**), and within a range of microtumor sizes, the model predicts mixtures of migratory and non-migratory microtumors as we actually observed experimentally for T47D microtumors (**Fig. 4G**) (Singh et al., 2016). Here, to reproduce our experimental results, we assumed that microtumor migration becomes visible when [sE-CAD] exceeds 80% of its steady state value with the corresponding time, T_migration_, while the exact value of this threshold does not affect our conclusions in this work. The network in **Fig. 4A** suggests that one can inhibit microtumor migration through inhibiting regulators (HIF-1α/MMPs/sE-CAD), while inter-tumor heterogeneity may complicate the outcomes. We present our modeling and experimental studies in the next sections.

### Anti-hypoxia treatment attenuates microtumor migration most effectively only when HIF-1α is inhibited at early time points

Based on our two-stage progression mechanism, we hypothesized that inhibition of HIF-1α is most effective during the initialization stage. **Figure 5A** shows simulated time courses of [sE-CAD] with no inhibitor (black line) and with HIF-1α inhibitor added at different times to 600 μm microtumors. The results show that a critical time exists (denoted as T_critical_) where, HIF-1α inhibition starting from T1 (T1 < T_critical_) can effectively prevent increase in [sE-CAD] (green line), thus inhibiting microtumor migration. In contrast, HIF-1α inhibitor treatment at T2 (T2 > T_critical_) cannot prevent further increase of already elevated [sE-CAD] (red line), hence the microtumor continues to migrate even in the presence of HIF-1α inhibitor. This implies that there is a treatment window (0 to T_critical_) for HIF-1α inhibitor to be effective in restricting microtumor migration. Due to dynamic heterogeneity, even microtumors of the same size have a distribution of T_critical_ (**Fig. 5B)**, which widens the boundary of this treatment window. Consequently, HIF-1α inhibition is effective in inhibiting migration in most of the microtumors when started as early as day 1, while some microtumors still migrate when the treatment is started on day 3 (T_critical_ < T < T_migration_). HIF-1α inhibition starting from day 4 or day 5 (T > T_critical_, T_migration_) is not effective, showing migration in most of the microtumors (**Fig. 5C-D**). This implies that it is too late to stop migration with HIF-1α inhibition when the bistable switch is committed to be turned on by the hypoxic microenvironment. Together, computational modeling strongly suggests that although the hypoxic microenvironment induces the migratory phenotype, HIF-1α inhibition is effective only for a small time window of treatment and the window boundary is blurred by the existence of dynamic tumor heterogeneity.

**Figure 5:**
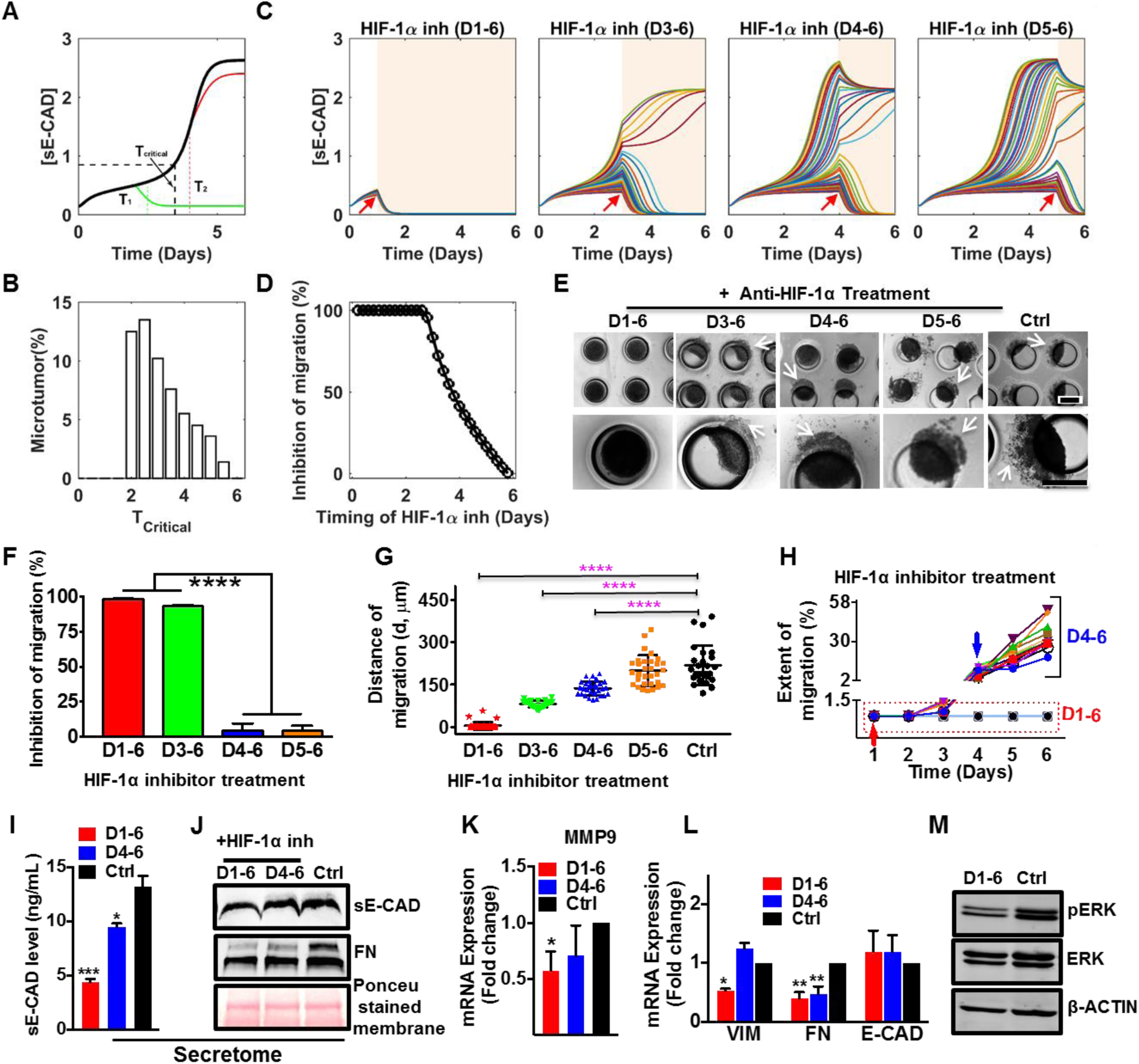
HIF-1α inhibitor treatment effectively attenuates microtumor migration only when HIF-1α is inhibited at early days of culture. (A) Mathematical model predicts a critical time (T_critical_, black dashed vertical line) to stop migration with HIF-1α inhibition for single microtumor. If the microtumor is treated with HIF-1α inhibitor before the critical time, the migration can be attenuated (green line). However, HIF-1α inhibitor has no effect on migration after the critical time (red line); (B) Distribution of the critical time, T_critical_ sampled with 1000 microtumors emphasizes inter-tumoral heterogeneity; (C) Dynamics of [sE-CAD] under HIF-1α inhibitor treatment started on day 1, day 3, day 4 and day 5, respectively. Fifty representative microtumors are shown for each case. The HIF-1α inhibitor treatment periods are shaded with light yellow background. HIF-1α inhibition at early time points effectively maintains [sE-CAD] levels below threshold; (D) HIF-1α inhibitor treatment (1.0 μM) started before day 3 inhibits microtumor migration significantly as suggested by simulation with 1000 large microtumors treated with HIF-1α inhibitor at different time points of culture; (E) Photomicrographs represent migratory status of 600 μm microtumors on day 6 with HIF-1α inhibitor treatment at different times. The effect of HIF-1α inhibition on microtumor migration was quantified by measuring percentage of microtumors showing inhibition of migration (F), distance of migration G) and extent of migration (H). The results demonstrated (F) significant inhibition of migration, (G) decrease in distance of migration at early treatments (D1-6 and D3-6); Each dot in G represents data for individual microtumor and each line in H represents migration kinetics for individual microtumor treated with HIF-1α inhibitor on day 1 (red arrow) or day 4 (blue arrow). HIF-1α inhibitor treatment started on day 1 downregulated (I) sE-CAD levels measured by ELISA and (J) sE-CAD and FN expression measured by western blot in conditioned media. HIF-1α inhibitor treatment (D1-6) also downregulated (K) *MMP9*; (L) *VIM and FN* mRNA expression; and (M) pERK protein expression in 600 μm microtumors. Data is presented as mean ±SEM. One way ANOVA followed by Tukey’ s test was used to analyze significance between different treatment groups. *p<0.05, **p<0.01, ****p< 0.0001 w.r.t. Ctrl (untreated) 600 μm microtumors.

To confirm our *in silico* results, we treated 600 μm microtumors with the chemical inhibitor of HIF-1α starting on day 1 (early stage), day 3, day 4 or day 5 (late stages) and continued until day 6 of the culture in each case. Interestingly, HIF-1α inhibition from day 1-6 or day 3-6 inhibited the microtumor migration significantly (only 3 and 11 out of 290 microtumors showed migration) (**Fig. 5E-H**). However, the HIF-1α inhibition after day 4 (D4-6, D5-6) did not block migration (distance of migration 137 ±25.15 μm and 199 ±54.44 μm, respectively) showing migratory behavior similar to untreated 600 μm microtumors (distance of migration 218.6 ±70.0 μm) (**Fig. 5E-H, Supplementary Fig. 5A-B)**. The distance and extent of migration showed tumor-to-tumor variation as shown by the distribution and the wider error bars in **Fig. 5G)** and this variation correlated with the delay in starting the HIF-1α inhibition treatment (**Fig. 5G-H**). More importantly, these migratory microtumors continued to migrate even in the presence of HIF-1α inhibitor (**Fig. 5H,** blue arrow**, Supplementary Fig. 5**).

To delineate the effects of HIF-1α inhibition on hypoxic secretome, we measured concentrations of sE-CAD by performing sandwich ELISA for sE-CAD in the CM. The results revealed that early inhibition of HIF-1α decreased the levels of sE-CAD by (66.76 ±3.27%) as compared to late inhibition (27.96±2.38%), (Fig. 5I). Additionally, FN and sE-CAD were measured in the CM of control and treated 600 μm microtumors by western blot. Early (day 1-6) HIF-1α inhibition decreased the levels of sE-CAD and FN in the CM (**Fig. 5J**) and significantly downregulated mRNA expression of *MMP9* (**Fig. 5K)** and *VIM* and *FN* (**Fig. 5L**), without significant effect on mRNA expression of the epithelial marker *E-CAD* compared to the untreated 600 μm microtumors. Early HIF-1α inhibition also significantly reduced pERK expression as compared to the untreated control (**Fig. 5M, Supplementary Fig. 5C**). Although HIF-1α inhibition on D4-6 reduced *FN* (both secreted in the CM and intracellular), this late HIF-1α inhibition could not reduce intracellular expression of *MMP9* and *VIM* mRNA (**Fig. 5K, L**). These observations are consistent with and support our mathematical model reinforcing positive feedback loop between sE-CAD and MMP9 in the maintenance stage, and suggest that once established, such positive feedback may contribute to the ineffectiveness of HIF-1α inhibition.

Together, these results suggest that early HIF-1α inhibition can prevent microtumor migration while late inhibition does not work effectively, thus confirming our computational predictions. The results also support our hypothesis that hypoxia is important only for the initiation stage of microtumor migration and provide a mechanistic explanation for the clinical failure of anti-hypoxia treatments (Patel and Sant, 2016).

### Blocking effects of sE-CAD by anti-sE-CAD antibody or preventing its shedding by MMP inhibitor effectively hinder microtumor migration

Here we tested the effect of temporal inhibition of sE-CAD on microtumor migration. Our simulations predict that early (day 1 or 3) inhibition of sE-CAD will prevent or to stop migration (**Fig. 6A**) by keeping [sE-CAD] at low level, so the bistable switch is not turned on to start the cascade necessary for microtumor migration. In the late phase (at or after day 4), although the sE-CAD level has exceeded the threshold causing some microtumors to migrate, the bistable switch is turned off with sE-CAD inhibition, and these microtumors cease to migrate further (**Fig. 6A-B**). That is, although a fraction of the migrated microtumor increases with the timing of the anti-sE-CAD **(Fig. 6B(i)),** the fraction of continuously migrating microtumor disappears no matter when the anti-sE-CAD is added **(Fig. 6B(ii)).** Thus, sE-CAD inhibition is effective in preventing microtumor migration independent of its treatment time and dynamic tumor heterogeneity. We obtained similar simulation results by inhibiting MMP-mediated sE-CAD shedding (**Supplementary Fig. 6A-B**). Together, these computational results suggest that unlike HIF-1α inhibition, the effectiveness of sE-CAD or MMP inhibition is independent of the time of treatment and tumor heterogeneity.

**Figure 6.**
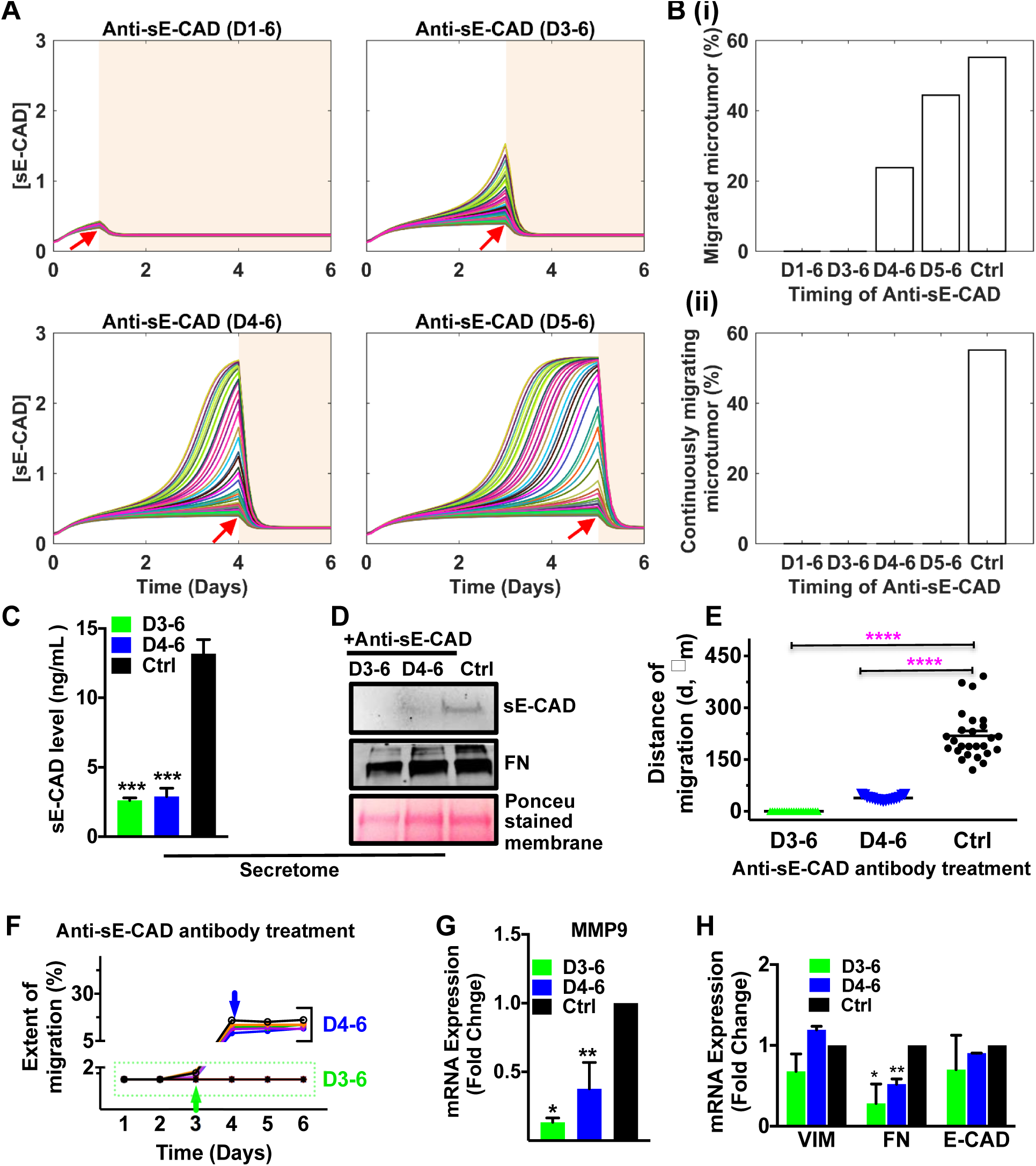
Inhibition of sE-CAD by antibody against ectodomain E-CAD attenuated collective migration in 600 μm microtumors. (A) The dynamics of [sE-CAD] for 50 microtumors post anti-sE-CAD antibody treatment started on day 1, day 3, day 4 and day 5 (red arrows); (B) The fraction of migrated microtumors (upper panel, i) and continuously migrating microtumors (lower panel, ii) with anti-sE-CAD treatment started on day 1, day 3, day 4 and day 5 (sampled with 1000 microtumors). To confirm results of mathematical modeling, 600 μm microtumors were treated with 40 μg/mL antibody against ectodomain E-CAD on day 3-6 and day 4-6. (C) Anti-sE-CAD antibody treatment significantly reduced sE-CAD levels measured by ELISA; (D) Western blot showed depletion of sE-CAD in conditioned media of 600 μm microtumors after anti-sE-CAD antibody treatment. However, secreted FN level was low only in day 3-6 treatment. Anti-sE-CAD treatment (green and blue arrows) inhibited migration in 600 μm microtumors represented as distance of migration; (E) and extent of migration (F); (G) Anti-sE-CAD antibody treatment significantly downregulated mRNA expression of *MMP9*; and (H) *FN* in 600 μm microtumors even after treatment starting on day 3 and day 4. Data are drawn from three independent experiments and are presented as means ±SEM. Significance between different treatment groups was analyzed by using one way ANOVA followed by Tukey’ s test. p*<0.05, **p<0.01, ***p<0.001 w.r.t. Ctrl (untreated) 600 μm microtumors.

To experimentally test the above model predictions, we blocked effects of sE-CAD in 600 μm microtumors using ecto-domain antibody (H-108, 40 μg/mL) against the extracellular domain of E-CAD, starting from day 3 and day 4 up to day 6 of culture. Anti-sE-CAD antibody treatment completely scavenged the sE-CAD and reduced the FN secreted in the CM as compared to the untreated control (**Fig. 6C, D**). As represented in the photomicrographs (**Supplementary Fig. 7A left panel**), day 3-6 anti-sE-CAD treatment completely inhibited collective migration. Although few microtumors treated with sE-CAD antibody on day 4-6 exhibited some migration (that was started before the treatment), the distance and extent of migration was significantly lower (38.4 ±6.5 μm and <25%, respectively) (**Fig. 6E, F**) compared to the distance and extent of migration (145.4 ±25.3 μm and 26-50%, respectively) achieved with HIF-1α inhibition on day 4-6 (**Fig. 5G-H**). Indeed, the single microtumor trajectories in **Fig. 6F** clearly showed that these already migrated microtumors stopped migration after the sE-CAD antibody treatment (blue arrow). The tight distribution of distance and extent of migration (**Fig. 6E, F**) also confirms our mathematical model predictions that all treated microtumors respond similarly with minimal effect on dynamic tumor heterogeneity. These findings were further confirmed by qRT-PCR results showing significant downregulation of *MMP9* (**Fig. 6G**) and mesenchymal marker, *FN* (**Fig. 6H**) with no significant change in *VIM* and *E-CAD* expression after sE-CAD antibody treatments on day 3 and day 4. We observed similar results in primary mBC microtumors where anti-sE-CAD antibody treatment (day 4-7) limited microtumor migration **(Supplementary Fig. 7B)** and downregulated *MMP9* mRNA expression **(Supplementary Fig. 7C)** and decreased sE-CAD levels **(Supplementary Fig. 7E)** but did not alter mesenchymal marker expression profiles **(Supplementary Fig. 7D)**. This could be due to already elevated mesenchymal marker expression in mBC cells derived from metastatic tumors **(Supplementary Figure 2)**.

**Figure 7.**
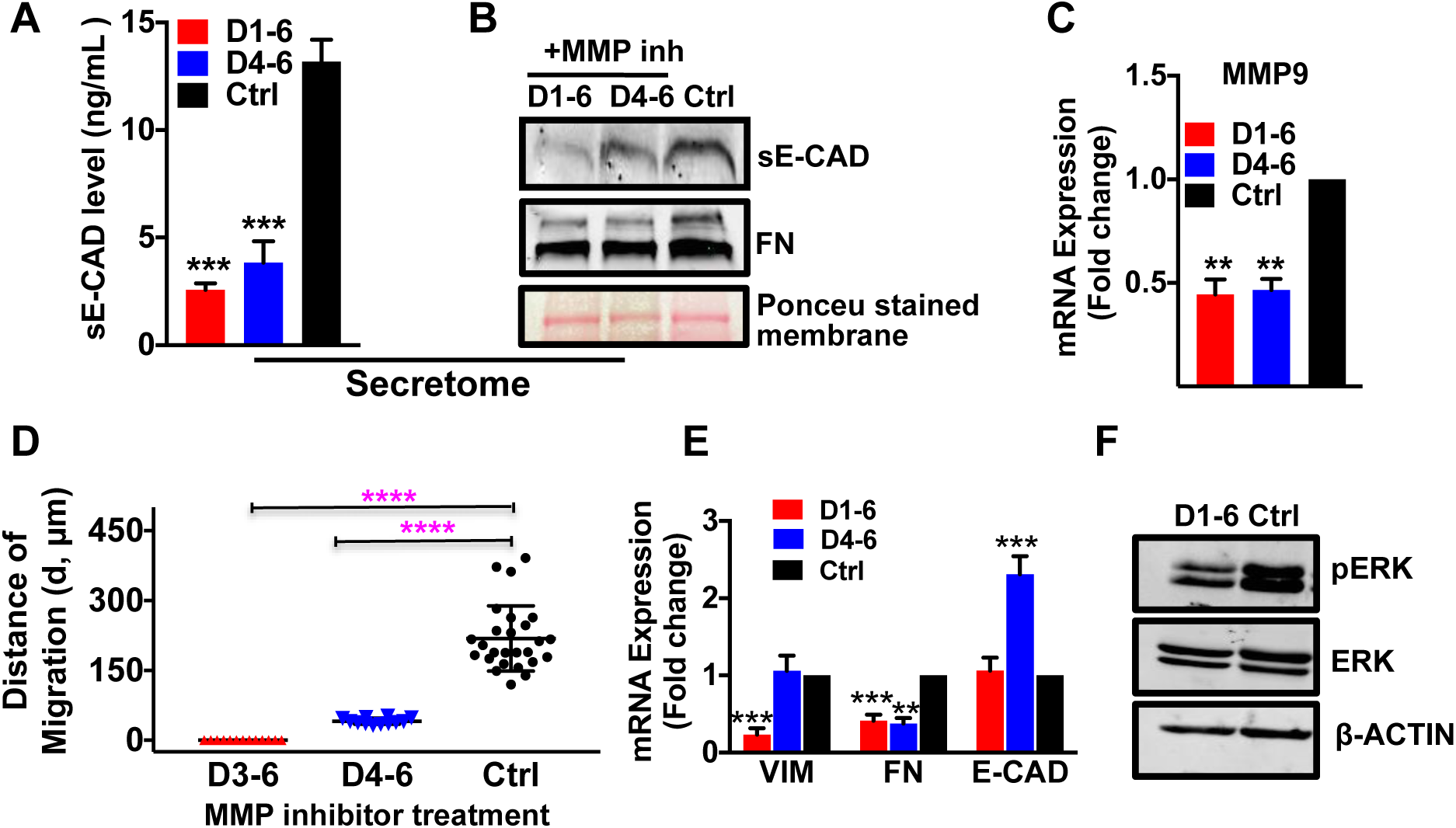
MMP inhibitor treatment attenuated collective migration in a time-independent manner. To investigate the effect of MMPs on dynamic regulation of sE-CAD levels in conditioned media and its effect on collective migration, 600 μm microtumors were treated with 20 μM of GM6001 (MMP inhibitor) on day 1-6 and day 4-6. (A) ELISA results showed decrease in sE-Cad levels in both the treatments as compared to Ctrl (untreated); (B) Western blot showing significantly reduced level of sE-CAD in the 600/CM of day 1-6 and day 4-6 inhibitor treated microtumors. Similarly, secreted FN level was significantly low in day 1-6 MMP inhibitor treatment; (C) MMP inhibitor treatment downregulated *MMP9* mRNA expression and (D) inhibited migration significantly; (E) MMP inhibition significantly downregulated *VIM* and *FN* mRNA expression; and (F) pERK in D1-6-treated 600 μm microtumors compared to untreated controls. Data is presented as means ±SEM from 3 independent experiments. One way ANOVA followed by Tukey’ s test was used to measure significance between treated groups and control. **p<0.01, ***p<0.001, ****p<0.0001; w.r.t. Ctrl (untreated) 600 μm microtumors.

Additionally, we blocked sE-CAD shedding in 600 μm microtumors through treatment with broad MMP inhibitor, GM6001 (20 μM) starting on day 1 or 4. MMP inhibition reduced sE-CAD shedding and level of secreted FN in the CM at both treatment times (**Fig. 7A, B)**. *MMP9* mRNA expression was significantly downregulated at both treatment time points (**Fig. 7C)**. We observed no migration in day 1-6 treatment whereas MMP inhibitor treatment starting from day 4-6 completely ceased the migration (**Fig. 7D**; **Supplementary Fig. 6C)**. MMP inhibition from day 1-6 significantly downregulated mRNA expression of *VIM,* and *FN* without significant effect on *E-CAD* expression compared to the untreated microtumors (**Fig. 7E).** Consistent with anti-sE-CAD antibody treatment, MMP inhibition on day 4-6 significantly downregulated *FN* expression without altering *VIM* mRNA expression. Similarly, MMP inhibition had a negative regulatory effect on ERK activation (**Fig. 7F**) as revealed by reduced pERK/ERK ratio (**Supplementary Fig. 6D**). It should be noted that *VIM* expression was downregulated only when HIF-1α and MMPs were inhibited early in the initialization stage while it had no effect when these molecules were inhibited in the maintenance stage. Similar to anti-sE-CAD antibody inhibition, MMP inhibition (day 4-7) prevented migration in primary mBC microtumors **(Supplementary Fig. 6E)**. qRT-PCR results showed downregulation of *MMP9* and *FN* along with decreased sE-CAD levels post inhibitor treatment as compared to untreated control **(Supplementary Fig. 6F-H)**.

Together, experimental results in T47D and multiple primary patient-derived microtumors confirmed the predictions from our mathematical model. MMPs, sE-CAD, and FN are the components of the positive feedback loop and are critical to the irreversible bistable switch, and to maintain the microtumor migration. HIF-1α, on the other hand, is a major trigger of the switch, which is dispensable once the switch is turned on. More importantly, these results confirmed our two-stage migration progression model that HIF-1α is important only in the initial stages of tumor progression. Consequently, inhibiting HIF-1α is only effective before T_critical_ that varies from tumor to tumor. Once MMPs, sE-CAD, and FN accumulate and exceed threshold values, targeting these molecules is more effective in inhibition of microtumor migration, and has less constrain on the treatment time and dynamic tumor heterogeneity.

## Discussion

Solid tumor growth generates multiple layers of cells that build a physical barrier to diffusion of oxygen, nutrients and metabolites (Vaupel and Harrison, 2004), leading to an hypoxic tumor microenvironment. This hypoxic microenvironment drives aggressive phenotype due to activation of various downstream signaling pathways (Nioka et al., 2013; Rankin and Giaccia, 2016; Singh et al., 2016; Vaapil et al., 2012). Hypoxic microenvironment further induces secretome containing sheddases and proteases that reshape the tumor microenvironment, modulate cellular phenotypes, and affect drug responses (Karagiannis et al., 2010; Miller et al., 2016; Miller et al., 2017; Rankin and Giaccia, 2016). These proteases are also involved in post-translational modification of membrane proteins, thus setting a cascade of events critical for motility and migratory behavior of tumor cells. One such product of these cleavages is sE-CAD, known to be cleaved by MMPs, secretases and a disintegrin and metalloprotease (ADAMs) to induce pro-inflammatory and pro-survival signals (Brouxhon et al., 2014b; Brouxhon et al., 2014c; Grabowska and Day, 2012; Hu et al., 2016). Various reports suggest that sE-CAD contributes to migratory phenotype in epithelial cells and helps in metastatic tumor progression (Brouxhon et al., 2014c; Hu et al., 2016; Patil et al., 2015). Ecto-domain E-CAD antibody prevents tumor cell migration and invasion by downregulating RTK, MAPK and PI3/Akt/mTOR signaling in squamous cell carcinoma and breast carcinoma (Brouxhon et al., 2014a; Brouxhon et al., 2013). Hypoxia-mediated increased secretion of FN and other extracellular matrix proteins is also known to help in tumor cell migration (Gilkes et al., 2014; Lu et al., 2012). Consistent with the literature, clinical data extracted using 3735 human breast tumor samples also showed higher levels of FN and MMP9 confirming the role of tumor secretome in tumor progression. We show that hypoxia and hypoxic secretome create a unique co-operative microenvironment that further activates complex cell signaling pathways promoting tumor migration. Therefore, it is desirable to have an experimental system that recapitulates this spontaneously formed hypoxic tumor microenvironment and the secretome because traditional 2D cell culture-based experimental systems are unable to capture this dynamic microenvironment.

To address these challenges, we engineered novel 3D microtumor model with controlled microenvironments to investigate the intricate core regulatory network in inducing migration. Indeed, we observed microtumor size-induced hypoxic signaling that leads to upregulation of mesenchymal markers and collective migration without loss of epithelial cadherin in large microtumors (Singh et al., 2016). The observed high levels of HIF-1α, MMPs, sE-CAD and FN in large microtumors suggest the interplay between hypoxia and hypoxic secretome for maintaining the migratory phenotype in large microtumors. Through theoretical analysis, we proposed an irreversible bistable switch mechanism by which tumor cells migrate collectively once the microtumor grows larger than a pre-set threshold. Our computational model successfully explained experimentally observed irreversible nature of the migratory phenotype in large microtumors and also predicted the effectiveness of temporal inhibition of HIF-1α, MMPs and sE-CAD on the microtumor migration. Most importantly, through computational and experimental approaches, we predicted and validated a novel mechanism of two-stage tumor progression where hypoxia is critical in initiating the migratory phenotype, which is then maintained by tumor secretome such as MMPs, sE-CAD and FN.

Many cancer treatments inevitably develop (chemo) resistance and tumor relapse, likely due to genetic and non-genetic heterogeneity and microenvironmental factors within a tumor (Easwaran et al., 2014; Huang, 2013; Koren and Bentires-Alj, 2015). Here, we demonstrated a dynamic heterogeneity of growth and migration at the level of individual microtumors, which couples to intra-tumor phenotypic heterogeneity at the cellular level evident by differential E-CAD and VIM protein expression. Different molecular species play temporally different roles in tumor progression, and proper choice of drug targets can minimize negative effects of dynamic heterogeneity on treatment efficacy (Donnenberg and Donnenberg, 2015b; Schmidt et al., 2016). Since the hypoxia-induced HIF-1α plays an important role only at the initial stage of activating the bistable switch, but not in the later maintenance stage of migration, HIF-1α inhibition can effectively halt tumor migration only if treated prior to the onset of migratory phenotypes, which is not easy in practice due to the stochastic dynamics of tumor growth. This further explains why HIF-1α inhibitor treatment failed to stop tumor progression in clinics, as it may be too late to stop the entire signaling cascade induced by HIF-1α in the already hypoxic tumors (Burroughs et al., 2013). Consequently, both our computational and experimental studies demonstrate that efficacy of HIF-1α inhibition to prevent tumor migration is significantly affected by treatment initiation time and tumor heterogeneity, which leads to variable treatment efficacy, consistent with the conflicting results observed in the clinical studies (Mack et al., 2008; Patel and Sant, 2016; von Pawel et al., 2000; Williamson et al., 2005).

Our two-stage progression mechanism suggests that the sE-CAD/MMP/FN axis mainly functions to maintain the migratory phenotype, and its inhibition is less affected by the stochastic tumor growth. Indeed, inhibition of sE-CAD shedding by MMP inhibitor as well as sequestering shed sE-CAD by ectodomain E-CAD antibody, both suppressed the collective migration in T47D and primary patient-derived microtumors as soon as the treatment was started. Unlike HIF-1α inhibition, sE-CAD inhibition was much less affected by tumor heterogeneity. Thus, our results support the two-stage tumor progression model and temporal dynamics of HIF-1α and sE-CAD. We propose sE-CAD as a novel potential target since there seem not to be an essential physiological role of sE-CAD (Grabowska and Day, 2012), pharmaceutical intervention to keep sE-CAD below a threshold level may not increase toxicities of concurrent anti-neoplastic regimens. Further, treatment strategies targeting inhibition of tumor growth, the proposed strategy here of targeting tumor secretome to constrain tumor migration may impose much less selection pressure as predicted from cancer evolutionary theory (Greaves and Maley, 2012).

Another important aspect of this study is our ability to cross-examine role of different regulators using complementary experimental systems such as pharmacological inhibition in hypoxic, migratory large microtumors and stimulation in non-hypoxic, non-migratory small microtumors of the same parent cells. For instance, late inhibition of sE-CAD by an ectodomain antibody stopped migration in large hypoxic microtumors by downregulating *FN* without significant effect on *VIM* expression. In contrast, stimulation with recombinant human sE-CAD proteins induced collective migration and *FN* upregulation in non-hypoxic, non-migratory small microtumors without any effect on *VIM* expression, suggesting that sE-CAD is not an effective regulator of *VIM*. Similarly, stimulation with hypoxic secretome (600/CM) upregulated *VIM* and *FN* in small microtumors, while early HIF-1α inhibition downregulated their expression in large microtumors, confirming the role of hypoxia in regulating *VIM* and *FN* expression. Together, these complimentary, yet controlled studies suggest that HIF-1α and sE-CAD regulate *FN* and *VIM* differentially. Our results also suggest a strong effect of temporal dynamics. Only early inhibition of microenvironmental factors (HIF-1, MMPs, and sE-CAD) could downregulate *VIM* expression while MMPs and sE-CAD inhibition in the later maintenance stage failed to reduce *VIM* expression, but could still prevent microtumor migration, suggesting that *VIM* is dispensable for migration. Thus, there may be requirement for other factors or simultaneous co-operation among multiple factors in the tumor microenvironment for tumor migration. For instance, in all the inhibition experiments (HIF-1, MMPs, and sE-CAD), migration was inhibited only if the treatment effectively inhibited MMP9 expression. These results are interesting since the role of epithelial and mesenchymal phenotypes in collective and single cell migration has been widely debated in recent years (Fischer et al., 2015; Nieto et al., 2016; Wong et al., 2014; Zheng et al., 2015). Our results suggest co-existence of multiple distinct phenotypes as indicated by the differential expression patterns of *VIM and FN*, and these phenotypes cannot be sufficiently characterized by individual mesenchymal marker as in the two controversial studies (Fischer et al., 2015; Zheng et al., 2015). Further studies are needed to examine whether these phenotypes actively contribute to migration or are simply passive outcomes of the microenvironment.

In summary, this study on microenvironmental changes in different stages of tumor progression provides an example of innovative ways to bypass major hurdles of tumor heterogeneity, drug resistance and drug-induced selection of more malignant cells in successful cancer treatments by targeting temporal dynamics of the evolving tumor microenvironments, resonant with other studies stressing the importance of dynamic pharmacology (Behar et al., 2013). While in this work we focused on the secretome factors, both our experimental system and computational approach can be readily applied to include couplings between the secretome network and intracellular signal transduction and gene regulation networks. The system can uniquely provide quantitative information of the temporal evolution of both, the microenvironment and cell phenotypes in an approachable experimental setting, and identify combined therapeutic strategies that overcome problems of drug resistance of existing ones (Donnenberg and Donnenberg, 2015b; Miller et al., 2017).

## Author Contributions

Conceptualization, SS and JX; Methodology, SS, JX, MS, VSD, AMW, SCW, and LPS; Clinical data analysis, JZ and JX; Computational modeling, JX, and XJT; Resources, SS, JX, VSD, SCW, and LPS; Data curation, SS, MS, JX, XJT, VSD, AMW, and SCW; Writing – Original Draft, SS, JX, MS, and XJT; Writing – Review and Editing, all authors; Supervision, SS and JX.

## Acknowledgments

We thank Dr. Wen Xie, Director, Center for Pharmacogenetics, University of Pittsburgh School of Pharmacy, for providing access to the core facilities. We also thank Drs. Barry Gold and Vinayak Sant, Department of Pharmaceutical Sciences, for critical reading and insightful suggestions on the manuscript. This work is supported by NIH (EB018575) and the Department of Pharmaceutical Sciences, School of Pharmacy, University of Pittsburgh to SS; the National Science Foundation (DMS-1462049) and the Pennsylvania Department of Health (SAP 4100062224) to JX; BC032981, BC044784, W81XWH-12-1-0415 and BC132245_W81XWH-14-0258 from the Department of Defense (VSD), the Hillman Foundation (VSD) and the Glimmer of Hope Foundation (VSD). The confocal microscope in the Center for Biologic Imaging at University of Pittsburgh is UPCI supported by 1S10OD019973-01 to SCW. Cytometry Facility is supported by CCSG P30CA047904.

## Supplemental Information

Supplementary Information includes supplemental experimental procedures, seven figures and two tables.

## Disclosure of Potential Conflicts of Interest

Authors declare no potential conflicts of interest.

## Materials and Methods

### Chemicals and Reagents

All chemicals and reagents were procured from Sigma-Aldrich (Milwaukee, USA) unless specified. All antibodies were purchased from Santa Cruz Biotechnology (California, USA) except anti-Vimentin (MA3-745), anti-beta actin (MA5-15739) from Thermo Scientific Inc., Rockford, USA; anti-ERK (9102S) and, anti-pERK (4370P) from Cell signaling Technologies Inc. (Massachusetts, USA); anti-HIF1-α (GTX127309) from GenTex, (California, USA) and anti-Fibronectin (610077) from BD Biosciences, California, USA). GM6001 (MMP inhibitor) was procured from Millipore (Ontario, Canada) and Methyl 3-[[2-[4-(2-adamantyl) phenoxy] acetyl]amino]-4-hydroxybenzoate (HIF-1α inhibitor) was obtained from Santa Cruz Biotechnology (California, USA). Human E-Cadherin Quantikine ELISA Kit was purchased from R&D Systems (Minneapolis, USA).

### Cell lines and cell culture

Breast cancer cell line T47D was purchased from American Type Culture Collection (ATCC) and cell culture supplies and media were obtained from Corning(®) and Mediatech(®), respectively unless specified. T47D cells were passaged and maintained in T75 flasks in Dulbecco Modified Eagle Medium (DMEM) (supplemented with 10% fetal bovine serum (FBS) (Hyclone, Utah, USA) and 1% penicillin-streptomycin in 5% CO_2_ at 37°C in a humidified incubator. Cell line authentication was done by University of Arizona, Genetics Core by using PowerPlex16HS PCR Kit. Briefly, genomic DNA was genotyped for 15 Autosomal STR loci and Amelogenin using Promega Power Plex16HS PCR kit and the electropherogram were analyzed using Soft Genetics, Gene Marker Software Version 1.85. Alleles were matched to STR profile and a minimum of 80% match threshold was used for standardization of STR Profiling and shared genetic history using ANSI database. Cells were maintained to attain 40-60% confluence for further seeding into hydrogel microwell devices.

### Primary cell cultures of malignant pleural effusions

Pleural effusions were collected from patients with metastatic breast cancer (mBC) or mesothelioma as waste materials during therapeutic drainage. Samples were anonymized by an honest broker and provided, along with relevant clinical information, to the laboratory according to University of Pittsburgh IRB exemption (0503126, VSD). The cells were cryopreserved in 1.25 mL vials (∼50 × 10^6^ cells/vial) in MEM containing 10% DMSO and 20% calf serum. The cryopreserved cells were carefully thawed in ice cold DMEM supplemented with 50% calf serum, resuspended in complete mammary epithelial growth medium (MEGM, Cat. No. CC-3151, Lonza, Walkersville MD) supplemented with 10% fetal bovine serum (Cat. No. SH30088.03, HyClone GE Healthcare Life Sciences) and 10% autologous cell-free pleural fluid. Complete MEGM consists of Epithelial Cell Basal Medium supplemented with bovine pituitary extract (BPE, Lonza Cat No. CC-4009G) human epidermal growth factor (rhEGF, 10 ng/mL, Lonza Cat. No. CC-4017G), hydrocortisone (0.5 μg/mL, Lonza Cat. No. CC-4031G), gentamicin sulfate plus amphotericin B (Lonza Cat. No. CC-4081G) and insulin (10 μg/mL, Lonza Cat. No. CC-4021G). The cells were initially plated in 10 cm petri dishes at a culture density of 1 - 1.5×10^5^ cells/cm^2^, grown to semi-confluence, trypsinized and split 1 to 3. All samples were analyzed at passage 0 and 1. Representative photomicrographs of these malignant pleural effusions used in this study are shown in **Supplementary Figure 2 and 3**.

### Multicolor flow cytometry on unpassaged metastatic breast cancer samples (MPE175)

Detailed methods used to disaggregate, stain and analyze unpassaged metastatic breast cancer samples have been described in detail (Donnenberg et al., 2010; Zimmerlin et al., 2011). Non-specific binding of fluorochrome-conjugated antibodies was minimized by pre-incubating pelleted cell suspensions for 5 min with neat decomplemented (56°C, 30 min) mouse serum (5μL) (Donnenberg and Donnenberg, 2015a; Donnenberg et al., 2013). The cells were then pelleted again by centrifugation and residual fluid aspirated. All antibodies were added at a fixed volume (2 μL) representing an approximate dilution of 1:5. Cells were first stained for surface markers. The order of addition was: CD326/EpCAM-APC (Miltenyi Cat. No. 130-098-118) CD45-APCCy7 (BD Biosciences, Cat. No. 348805), CD44-PECy7 (Abcam, Cambridge, MA, Cat. No. ab46793), CD90-APC (BD Biosciences, San Jose CA, Cat. No. 559869), CD14-PECy5 (Beckman-Coulter, Brea CA, Cat. No. IM2640U), CD33-PECy5 (Beckman-Coulter, Cat. No. IM2647U) and Glycophorin A-PECy5 (BD Biosciences, Cat.No.559944), CD73-PE (BD Pharmingen, Cat. No. 550257). Unbound antibodies were washed and cells were fixed with 2% methanol-free formaldehyde (Polysciences, Warrington, PA). Cells were then permeabilized with 0.1% saponin (Beckman Coulter) in phosphate buffered saline with 0.5% human serum albumin (10 min at room temperature); cell pellets were incubated with 5 μL of neat mouse serum for 5 min, centrifuged and decanted. The cell pellet was disaggregated and incubated with 2 μL of anti-pan cytokeratin (CTK, Abcam, Cat. No. ab52460) and vimentin-(VIM, BD Pharmingen Cat. No. 562338) for 30 min. Cells pellets were diluted to a concentration of 10 million cells/400 μL of staining buffer and DAPI (Life Technologies, Grand Island NY, Cat. D1306) was added 10 min before sample acquisition, to a final concentration of 7.7 μg/mL and 40 μL/10^6^ cells (Donnenberg et al., 2007). In this application, DAPI was used as a DNA stain in fixed permeabilized cells because it permits removal of artifacts associated with tissue digestion and also provides information on cell DNA content (Donnenberg and Donnenberg, 2015a). Stained cells were acquired on a Beckman Coulter Gallios cytometer calibrated to pre-established PMT target channels. Offline compensation and analyses were performed using VenturiOne software (Applied Cytometry, Dinnington, Sheffield, UK) as detailed previously (Donnenberg et al., 2013). Spectral compensation matrices were calculated for each staining combination within each experiment using single-stained mouse IgG capture beads (Becton Dickinson, Cat. No. 552843) for each tandem fluorochrome, and BD Calibrite beads for PE and APC controls (Becton Dickinson, Cat. No. 340486).

### 3D culture and microtumor fabrication

Non-adhesive hydrogel microwell devices of 150 and 600 μm were microfabricated using polyethylene glycol dimethacrylate (PEGDMA, 1000Da) and polydimethyl siloxane (PDMS) molds as described previously (Singh et al., 2015). PDMS molds with 150 and 600 μm posts (1:1 aspect ratio of height: diameter) were placed on PEGDMA solution (20% w/v) mixed with photoinitiator (Irgacure-1959, 1% w/v, Ciba AG CH-4002, Basel, Switzerland) and photo-crosslinked using the OmniCure S2000 curing station (EXFO, Mississauga, Canada). Subsequently, T47D cell suspension in growth media (200 μL, 4.0 × 10^6^ cells/device) was seeded on the 2×2 cm^2^ hydrogel microwell device and cultured in a humidified 5% CO_2_ incubator. Microtumors were cultured up to 6 days and harvested on day 6 and processed further as required or frozen at -80°C. Methods for isolation of primary cells from primary metastatic breast cancer (mBC), mesothelioma and patient-derived mouse xenograft (PDX) as well as fabrication of 3D microtumors from these primary tumor cells and xenografts is described in **Supplementary Information** (Materials and Methods).

### Clinical data for FN and MMP9 expression in patient tumors

For FN1 and MMP9 expression profile analysis in patient tumors, we integrated high quality metadata of human that were obtained by AffyU133Plus2 platform expression arrays from public repositories, such as GEO. 3735 Homo sapiens samples were included in the results. The original targeted gene expression profile in nine subcategories of cancers as well as normal tissue was extracted and normalized by Genevestigator®.

### Treatment of microtumors with conditioned media (CM) and various inhibitors

To study the effect of hypoxic secretome, 150 μm microtumors were treated with conditioned media of 600 μm microtumors (denoted as “600/CM”) from day 3 to day 6. To investigate the role of soluble E-CAD (sE-CAD) on microtumor migration, 150 μm microtumors were treated with recombinant human E-CAD, Fc chimera protein (20 μg/mL, R&D Systems, Minneapolis, USA) for pre-determined time.

HIF-1α was inhibited chemically by treatment of 600 μm microtumors with growth media containing HIF-1α inhibitor, methyl 3-[[2-[4-(2-adamantyl) phenoxy] acetyl] amino]-4-hydroxybenzoate (1.0 μM) at different time points. MMP inhibition was achieved by treating the 600 μm microtumors with of broad spectrum MMP inhibitor (GM6001, 20 μM). To scavenge sE-CAD secreted in the conditioned media, 600 μm microtumors were treated with ectodomain antibody (H-108, 40 μg/mL, Santa Cruz Biotechnology) in the growth media from day 3 to day 6 or day 4 to day 6.

After CM and various inhibitor treatment studies, mRNA were isolated from control (untreated) and treated microtumors and EMT marker expression was determined by qRT-PCR. Similarly, protein expression was estimated by western blotting.

### Kinetics of microtumor migration

The aggressive phenotype of microtumors was assessed by measuring the distance of migration (d) and extent of migration (%), respectively. The distance of migration (d) was estimated by measuring the average length of straight-lines drawn perpendicular to periphery of the well up to the leading edge, covering the entire migratory front using ImageJ. Of note, distance of migration for each microtumor was average of approximately ≥ 10 such perpendicular lines and was represented as the distance in microns. The extent of migration is the percentage of the area of the microtumor that is migrated out of the microwells (X) normalized with the total area of the microtumor (Y), [extent = X/Y *100]; see inset in **Fig. 1B**) and was measured by ImageJ using analyze > set scale> Ctrl+T> measure tools.

### Protein extraction and western blotting

Frozen microtumor samples were lysed in radioimmunoprecipitation assay (RIPA) buffer containing Tris (50 mM, pH 7.4), sodium chloride (150 mM), sodium deoxycholate (0.5%), NP-40 (1%), phenylmethane sulfonyl fluoride (0.05 mM), and protease inhibitor cocktail for mammalian tissue extract and phosphatase inhibitor (Sigma-Aldrich Inc., Milwaukee, USA). Proteins secreted in the conditioned media (sE-CAD and FN) were measured by resolving 50 μL of conditioned media on 8-10 % polyacrylamide gel, transferring them to PVDF membrane and probing them with respective antibodies. Of note, CM of 150 and 600 μm microtumors were collected from the hydrogel microwell devices that have same number of cells. To match the cell number of 150 and 600 μm microwell devices, cells/microtumor was calculated for both 150 and 600 μm microtumors. Subsequently, the dimensions of each device were calculated for bearing the same cell number. Western blot was performed for primary antibodies against Vimentin (1:1000), Fibronectin (1:1000), E-CAD (1:1000), pERK (1:1000), ERK (1:1000), and β-actin (1:2000) as described previously (Singh et al., 2016). Membranes were scanned by Odyssey Infrared Imaging System (LI-COR® Biosciences, Lincoln, USA) and colored images were converted to grayscale images by using Odyssey 3.0 software. Quantification of blots was done by ImageJ Software (National Institute of Health, USA) and normalization of densitometry was done by comparing with β-actin. The data are expressed as fold change compared to control/600 (n=3).

### Measurement of shed sE-CAD in the conditioned media by ELISA

sE-CAD levels in the conditioned media (CM) of 150 and 600 μm microtumors and inhibition studies were measured by quantitative ELISA (R&D Systems, DCADE0) according to the manufacture’ s protocol. All the CM were diluted (1:2) using dilution buffer from the kit before assay as described in the instruction manual. The standards and samples (50 μL) were pipetted into the E-CAD antibody pre-coated wells of microplate and incubated for 2 h at room temperature on rocking platform. Any unbound material was removed by four times washing with wash buffer, an enzyme-linked polyclonal antibody specific for E-CAD was added to the wells and incubated for 2 h at room temperature on rocker. Any unbound antibody-enzyme reagent was washed with wash buffer, followed by addition of a substrate solution and incubation for 30 min to develop the color. The reaction was stopped using stop solution and absorbance was measure at 450 nm. Each sample was measured in duplicate. The concentration of sE-CAD were determined by extrapolating the values on the standard curve and multiplied by dilution factor (2) and represented as ng/mL.

### Immunostaining and confocal microscopy

E-Cadherin-GFP expressing microtumors were cleared using CUBIC R1 (Susaki et al., 2014). The clearing solution was removed by washing with CUBIC IHC buffer (phosphate buffered saline, PBS; 0.1% Triton X-100; 0.5% bovine serum albumin, BSA; 0.01% Sodium Azide) and then stained with anti-Vimentin (Abcam, ab73159) for 24 h. Samples were then washed and incubated with anti-chicken-Cy5 for 24 h (Jackson ImmunoResearch, 703-175-155). Samples were washed again and placed in CUBIC R2 without triethanolamine (Watson et al., 2017) containing Hoechst for 24 h prior to imaging. All incubation steps were done at 37°C with gentle movement. The immunostaining protocol was optimized to ensure antibody penetration throughout the volume of microtumors. Volumes were acquired over whole microtumors using a Nikon Ti-E equipped with a Bruker SFC and excitation at 405, 488 and 640 nm. The Voxel size was 0.4 × 0.4 × 5.0 μm using an Apo 40x WI λS DIC N2 objective. Images were prepared in NIS Elements v4.51.00. Preparation of publication quality images included rolling ball background correction and local contrast enhancement.

### RNA isolation and qRT-PCR analysis

RNA was isolated from all the samples by using GeneJET RNA purification kit (Thermo Scientific, Lithuania, EU) as per manufacturer’ s protocol. RNA quantity was measured by absorbance ratio at 260/280 nm and integrity was verified on 1% agarose gel. mRNA expression of EMT markers such as Vimentin (*VIM*), Fibronectin (*FN*) and E-cadherin (*E-CAD*) and matrix metalloprotease 9 (*MMP9*) was analyzed by qRT-PCR using iTaq Universal SYBR Green One-Step RT-PCR kit (BioRad Laboratories Inc., USA) and 7500 Fast Real-Time PCR System (Applied Biosystems, California, USA) using β-Actin as control house-keeping gene. The mRNA expression was quantified using the 2^-ΔΔCt^ method and presented as the mean ±SEM as fold change compared to controls (150 or 600 μm microtumors). Primer sequences are given in Supplementary Table 1.

### Gelatin zymography

MMPs are responsible for cleavage of ectodomain E-CAD to generate 80 kDa sE-CAD. To estimate the levels of MMPs in the CM, gelatin zymography was performed as described by Frankowski et al. (Frankowski et al., 2012). Briefly, 50 μL of CM was mixed with non-reducing sample buffer and separated on 7% polyacrylamide gel containing 1.0% porcine gelatin Type A (Sigma). Gel was washed thrice using wash buffer containing 25% v/v Triton X-100. Subsequently, the gel was incubated in developing buffer containing 50 mM Tris, 10 mM CaCl2 and 0.02% sodium azide, for 48 h at 37°C. Staining was done for 40 min using Coomassie Blue staining solution and destaining was done until the bands appeared. Imaging was done using GelDoc (BioRad) and quantification was done by densitometric analysis of bands by ImageJ.

### Statistical analysis

Statistical analysis was done using GraphPad Prism7. All the values were presented as mean ±SEM. Unpaired t-test was used to compare the significance between two groups. For multiple comparisons (effects of inhibitor treatments), data were analyzed by one-way analysis of variance (ANOVA) followed by a Tukey’ s test. A *p*-value less than 0.05 was considered significant.

### Mathematical modeling and analysis

A set of ordinary differential equations (ODEs) to model the N-terminal E-cadherin shedding is applied in the theoretical analysis (details can found in the Supplementary material). The model quantitatively captures the dynamics of the core regulatory network that governs the N terminal E-cadherin shedding in hypoxic microenvironment. The default state is the non-migratory phenotype, in which the levels of the HIF-1α, MMPs and sE-CAD are low. The quantitative model consisted of 4 ODEs with 20 kinetic rate parameters (Supplementary Table 2). The bifurcation analysis was performed with the software Oscill8 (http://oscill8.sourceforge.net/). For HIF-1α inhibition simulation, the HIF1-α-mediated expression rate of MMP is set to 1/10 of its original value, while for the sE-CAD inhibition the sE-CAD-mediated expression rate of ERK is set to 1/10 of its original value. For the MMP inhibition, the shedding rate of sE-CAD mediated by MMPs is set to 1/10 of its original value. All the concentrations are in a reduced unit. More detail is provided in SI materials and methods.

